# Stopping Transformed Growth with Cytoskeletal Proteins: Turning a Devil into an Angel

**DOI:** 10.1101/221176

**Authors:** Bo Yang, Haguy Wolfenson, Naotaka Nakazawa, Shuaimin Liu, Junqiang Hu, Michael P. Sheetz

## Abstract

The major hallmark of cancer cells is uncontrollable growth on soft matrices (transformed growth), which indicates that they have lost the ability to properly sense the rigidity of their surroundings. Recent studies of fibroblasts show that local contractions by cytoskeletal rigidity sensor units block growth on soft surfaces and their depletion causes transformed growth. The contractile system involves many cytoskeletal proteins that must be correctly assembled for proper rigidity sensing. We tested the hypothesis that cancer cells lack rigidity sensing due to their inability to assemble contractile units because of altered cytoskeletal protein levels. In four widely different cancers, there were over ten-fold fewer rigidity-sensing contractions compared with normal fibroblasts. Restoring normal levels of cytoskeletal proteins restored rigidity sensing and rigidity-dependent growth in transformed cells. Most commonly, this involved restoring balanced levels of the tropomyosins 2.1 (often depleted by miR-21) and 3 (often overexpressed). Restored cells could be transformed again by depleting other cytoskeletal proteins including myosin IIA. Thus, the depletion of rigidity sensing modules enables growth on soft surfaces and many different perturbations of cytoskeletal proteins can disrupt rigidity sensing thereby causing transformed growth of cancer cells.

## Introduction

For normal cell survival, complex cellular mechanical functions need to sense the microenvironment and develop the proper signals for growth. When cells encounter the wrong environment, the output from these sensing events will activate cell death. Matrix rigidity is one of the most critical aspects of the microenvironment for normal development and regeneration, but it is essentially ignored by cancer cells since they grow on very soft surfaces. This is the basis for the soft agar assay, which is a standard test for the malignancy level of cancers (Hamburger and Salmon, 1977).

We recently described the rigidity sensing apparatus as a cytoskeletal protein complex that contracts matrix to a fixed distance; and if the force generated by this contraction exceeds about 25 pN, the matrix is considered rigid (Wolfenson et al., 2016). This is just one of a number of modular machines that perform important tasks in cells, including, e.g., the clathrin-dependent endocytosis complex (McMahon and Boucrot, 2011). Such machines typically assemble rapidly from mobile components, perform the desired task, and disassemble in a matter of seconds to minutes. They are activated by one set of signals and are designed to generate another set of signals. The cell rigidity sensing complex is a 2-3 μm-sized modular machine that forms at the cell periphery during early contact with matrix (Ghassemi et al., 2012; Meacci et al., 2016; Saxena et al., 2017; Wolfenson et al., 2016; Yang et al., 2016). It is powered by sarcomere-like contractile units (CUs) that contain myosin-IIA, actin filaments, tropomyosin 2.1 (Tpm 2.1), α-actinin 4, and other cytoskeletal proteins (Meacci et al., 2016). The number of CUs depends upon EGFR or HER2 activity as well as substrate rigidity (Saxena et al., 2017). Further, the correct length and duration of contractions are controlled by receptor tyrosine kinases through interactions with cytoskeletal proteins (Yang et al., 2016). CUs are activated in spreading cells, and on rigid surfaces they stimulate the formation of mature adhesions. However, on soft surfaces, contractions are very short-lived and adhesions rapidly disassemble, leading to cell death by *anoikis* (Meacci et al., 2016; Wolfenson et al., 2016). Since cancer cells fail to activate *anoikis* pathways on soft matrices, and since rigidity-sensing CUs can block growth on soft surfaces, we postulated that CU depletion in cancer cells can enable transformed growth on soft agar.

Cytoskeletal proteins are integrated into many complex cellular functions and their roles are well studied in normal cells (Fletcher and Mullins, 2010). However, the role of cytoskeletal components in cell transformation and cancer development is still not clear. Mutations and abnormal expression of various cytoskeletal or cytoskeletal-associated proteins have been reported in many cancer studies (Fife et al., 2014): Myosin IIA has been identified as a tumor suppressor in multiple carcinomas (Schramek et al., 2014) (Conti et al., 2015); The expression level of Tpm 2.1 is highly suppressed in a variety of cancer cell lines (Raval et al., 2003); Tpm 3 (including Tpm 3.1 and Tpm 3.2), another tropomyosin isoform, is the predominant tropomyosin in primary tumors and tumor cell lines (Stehn et al., 2013); and α-actinin 4 is reported to be a tumor suppressor in certain cases (Menez et al., 2004; Nikolopoulos et al., 2000) but an activator in others (An et al., 2016). Although many cytoskeletal proteins correlate with either tumor suppression or activation, it is not clear how they could be involved in cell growth control related to cancer.

A potential relation between cancer and the function of cytoskeletal proteins was suggested by studies of the rigidity sensing basic mechanisms. In our recent studies we found that rigidity sensing activity was missing in MDA-MB-231 cancer cells but was normal in the control MCF 10A epithelial cells as defined by local contractions of sub-micrometer pillars (Wolfenson et al., 2016). In contrast, both cell lines developed actin flow-driven traction forces on the substrates. Previous studies showed that restoration of normal levels of Tpm 2.1 blocked transformed growth of MDA-MB-231(Bharadwaj et al., 2005; Raval et al., 2003) and metastasis (Zhu et al., 2008), which correlated with the restoration of rigidity sensing after Tpm 2.1 expression (Wolfenson et al., 2016). Conversely, depletion of Tpm 2.1 in the normal MCF 10A cells caused transformation as well as disruption of rigidity sensor formation (Raval et al., 2003; Wolfenson et al., 2016). Further, removing another CU component, α-actinin 4, in mouse embryonic fibroblasts (MEF), disrupted rigidity sensors and enabled rapid growth on soft matrices (Meacci et al., 2016). These findings indicated that not only were alteration of cytoskeletal protein levels involved in transformation but also there was a strong correlation between the loss of rigidity sensing and transformation.

Here we report that widely different transformed cancer cells lack rigidity sensing and can be restored to a rigidity-dependent growth state by restoring rigidity sensing. In correlation with the restoration of rigidity sensing, the cells undergo a state change from transformed to normal that correlates with many differences previously observed between those two states, including changes in contractile behavior and adhesion and cytoskeletal organization. Different cytoskeletal proteins can be altered to reversibly cause the transition between the normal and transformed states. This all indicates that the transformed state is the default state that is activated when cells are unable to sense matrix rigidity and that cancer cells are stably in that state because of specific cytoskeletal protein alterations.

## Results

### Fundamental Differences between Normal and Transformed Cancer Cells

To understand the nature of the differences between normal fibroblasts and transformed cancer cells, we analyzed the forces that normal and cancer cells develop on discreet adhesion sites (as measured with 500 nm diameter pillars) because cell adhesions are similar on continuous surfaces and on 500 nm pillars coated with fibronectin (Ghassemi et al., 2012). In the case of HFF (human foreskin fibroblasts) cells, they formed 2-3 μm-sized contractile units (CU) (Figure 1 A blue color vectors) at the cell edge that tested substrate rigidity. A typical CU contained a pair of pillars that were contracted toward each other by an average of 60 nm with a half deflection time of 30 s (Figure 1 B). More than 25% of the pillars pulled by HFF cells during spreading contributed to CU formation (Figure 1 E). When pillars were part of contractile pairs, they had a narrower distribution of displacements than pillars that were pulled centripetally by cells (Figure 1 C). Further, the average displacement of the contractile pairs was significantly greater than the radial contractions (Figure 1 D). In contrast, HT1080 cancer cells (a fibrosarcoma line from an untreated patient that carried an IDH1 mutation (Rasheed et al., 1974)), had very few contractile pairs and the radial contractions were significantly greater in magnitude than those in the HFFs (Figure 1 H, I and J). There was no significant difference between the total cell spreading areas of HFF and HT1080 cells after 20 minutes seeding on pillars (Figure 1 F and G). Thus, overall patterns of contractility were dramatically different between normal fibroblasts and the HT1080 fibrosarcoma cells, indicating that they were in fundamentally different cell states.

**Figure 1.**
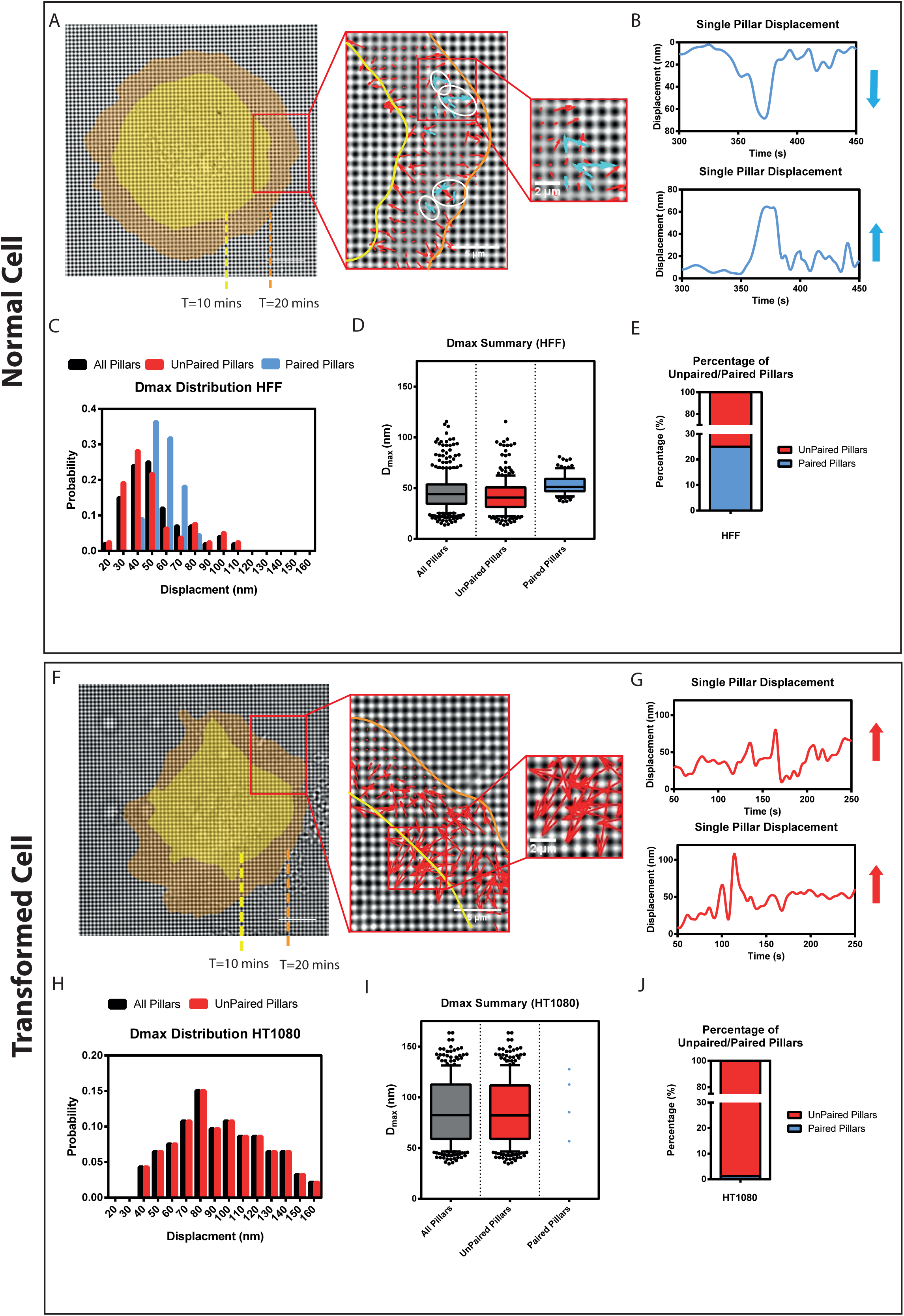
Fundamental differences between normal and transformed cells during initial spreading. (A and F) Actual CUs observed with a HFF cell (A) or a HT1080 cell (F) spreading on FN-coated rigid pillars (k=8.4 pN/nm). Arrows represented pillar movement: Blue, detected CUs; Red, non-CUs. The white circles marked several examples of identified CUs. Cell edges were marked in yellow or orange corresponding to time points 10 or 20 mins after seeding, respectively. (Scale bar is 10 μm) (B and G) Displacement vs. time plots of two nearby pillars that were part of a CU in HFF (B) or HT1080 (G) cells. Blue (HFF) or Red (HT1080) vectors represent the relative displacement directions and magnitude of those two pillars. (C and H) Histogram distribution of pillar maximum displacements from one HFF (C) or one HT1080 (H) cell. Black represented all tracked pillars; Red represented pillars that did not form any CUs during imaging; Blue represented pillars that formed CUs. (>100 pillars were analyzed) (D and I) Box and whiskers plots of pillar maximum displacement summary from 3 HFF (D) or 3 HT1080 (I) cells. (>300 pillars were analyzed) (E) Bar graph of unpaired/paired pillar displacement percentages of HFF (E) or HT1080 (J) cells. (>300 pillars from 3 different cells were analyzed)

### Transformed Cells Lack Local Contractions

To further examine the relationship between rigidity sensing CU formation and transformed growth in different cell backgrounds, we tested three other cancer lines from different organs and a non-cancerous transformed cell line, each randomly selected. The bases of transformation for these cells were all different: 1) Cos7 cells were derived from African green monkey kidney fibroblast cells by SV40 transformation; 2) MDA-MB-231 was a human breast cancer line that formed tumors in nude mice (Freedman and Shin, 1974; Hollestelle et al., 2007); 3) SKOV3 was a human ovarian adenocarcinoma line with an epithelial-like morphology (Fogh et al., 1977); 4) LLC was a lung carcinoma line from a C57BL mouse (Mayo, 1972). CU activity was measured as previously described (Saxena et al., 2017) and then normalized to cell spreading area. Consistent with previous publications, HFF cells generated 39 CUs/100 μm^2^ on rigid pillars (k=8.4 pN/nm) and 24 CUs/100 μm^2^ on soft pillars (k=1.6 pN/nm) per 10 minutes during initial spreading (Figure 2 A and B). In contrast, all the transformed cells produced less than 2 CUs/100 μm^2^ on the two pillar types (Figure 2 A and B). To test whether cells were able to distinguish between rigid and soft surfaces at a later spreading stage, we plated the 6 different cell lines on stiff (2 MPa) and compliant (5 kPa) fibronectin-coated PDMS surfaces for 6 hours. Cells were then fixed and stained with paxillin and actin as markers for adhesion formation and morphology change, respectively (Figure 2 C). As previously described (Prager-Khoutorsky et al., 2011), HFF cells polarized on rigid PDMS and spread less on soft PDMS in a round shape. However, all five transformed cell lines showed no significant difference in cell polarization level or adhesion size on the surfaces with a 400-fold difference in rigidity (Figure 2 D and E). Since transformation was defined classically as growth on soft agar (Hamburger and Salmon, 1977), we cultured the different cell lines in soft agar for 7 days. All five transformed cell lines formed colonies while HFF cells barely survived (Figure 2 F and G). Thus, none of the transformed lines developed a significant number of CUs for rigidity sensing, and this was consistent with their inability to react to differences in matrix rigidity and hence their ability to grow on soft surfaces.

**Figure 2.**
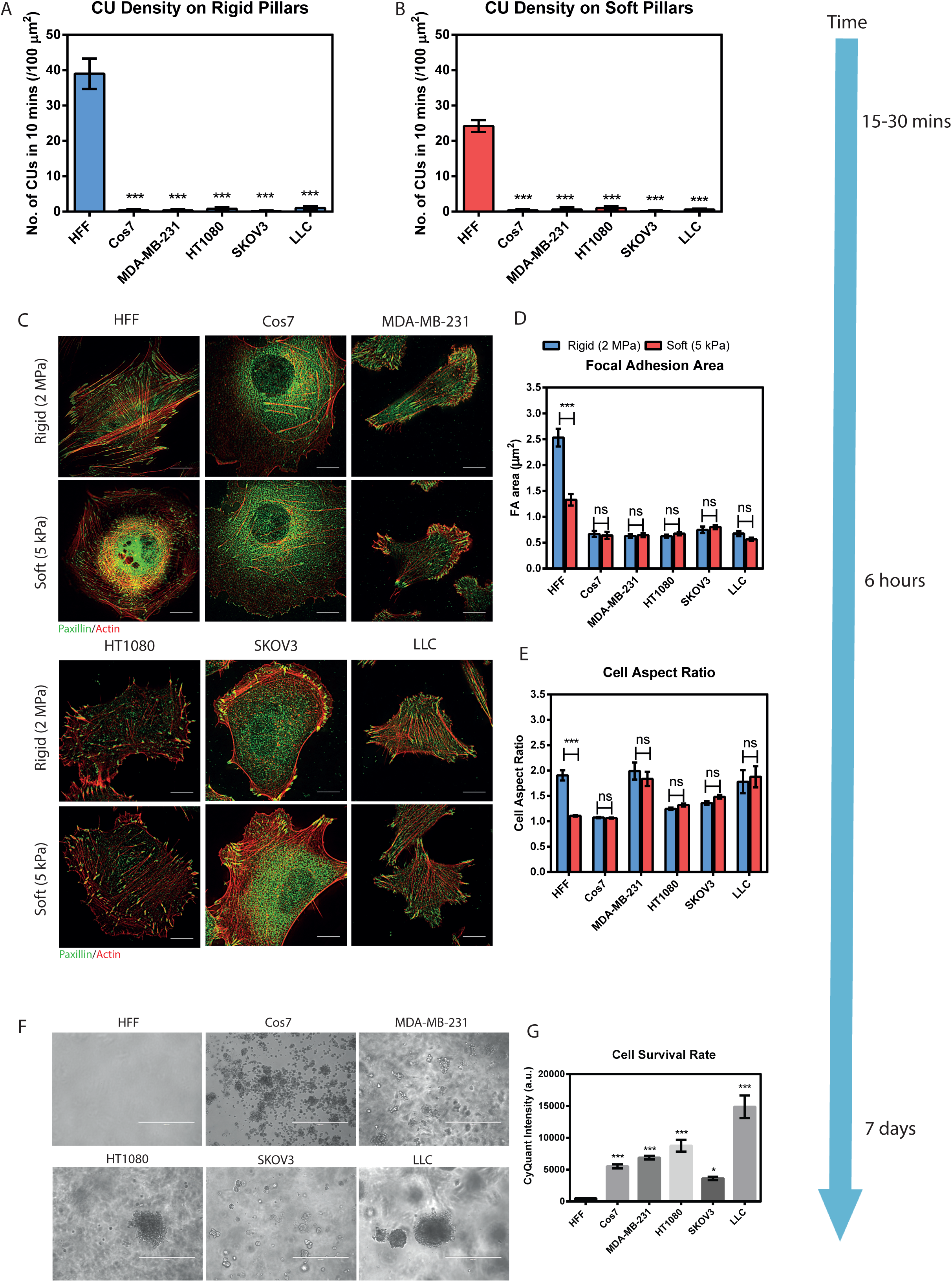
Transformed cells lack rigidity sensing. (A and B) Average number of CUs/ 100 μm^2^ generated by HFF or various transformed cell lines on rigid (k=8.4 pN/nm) and soft (k=1.6 pN/nm) pillars per 10 minutes, respectively. (C) Staining for actin (red) and paxillin (green) in HFFs and other transformed cells on hard (2 MPa) and soft (5 kPa) PDMS surfaces after 6 hours plating. (Scale bar is 10 μm) (D and E) Mean single focal adhesion (FA) area (D) and cell aspect ratio (E) of HFFs and various transformed cells after 6 hours plating on soft (red) and rigid (blue) PDMS surfaces. (F and G) Soft agar assay showing growth of various transformed cells but not HFF cells after 7 days culturing. (Scale bar is 400 μm) Cell proliferation rate was analyzed by measuring CyQuant intensity. (Error bars represented SEMs. Experiments were repeated >3 times; *** stands for p<0.001; ** stands for p<0.01; * stands for p<0.05)

### Transformed Cells Lack Rigidity Sensing Activity due to Altered Expression Levels of Contractile Unit (CU) Components

To determine if the transformed cells had altered protein levels in the rigidity sensor components, we performed a western blot screening of the known CU proteins (Figure 3 A), including kinases (EGFR, HER2, and ROR2) and cytoskeletal proteins (Myosin IIA, Tpm 2.1 and Tpm 3) (Figure 3 B). Interestingly, the 5 different transformed cell lines lacked at least one CU component. In addition, the expression levels of selected cytoskeletal proteins followed a similar trend in different transformed backgrounds. Both myosin IIA and Tpm 2.1 levels were suppressed, while Tpm 3 (Tpm 3.1 and 3.2) expression level was increased in most cancer lines (Figure 3 B and Supplementary Figure 2). On the other hand, the pattern of RTK levels varied from cell line to cell line (Figure 3 B and Supplementary Figure 2). Taken together, we observed different patterns of protein depletion in transformed cells, including cytoskeletal components and receptor tyrosine kinase levels. This raised the questions: 1) Could CUs be restored by restoring normal levels of only the altered cytoskeletal components? 2) Would alteration of another cytoskeletal protein in restored cells inhibit CU formation and again cause transformed growth?

**Figure 3.**
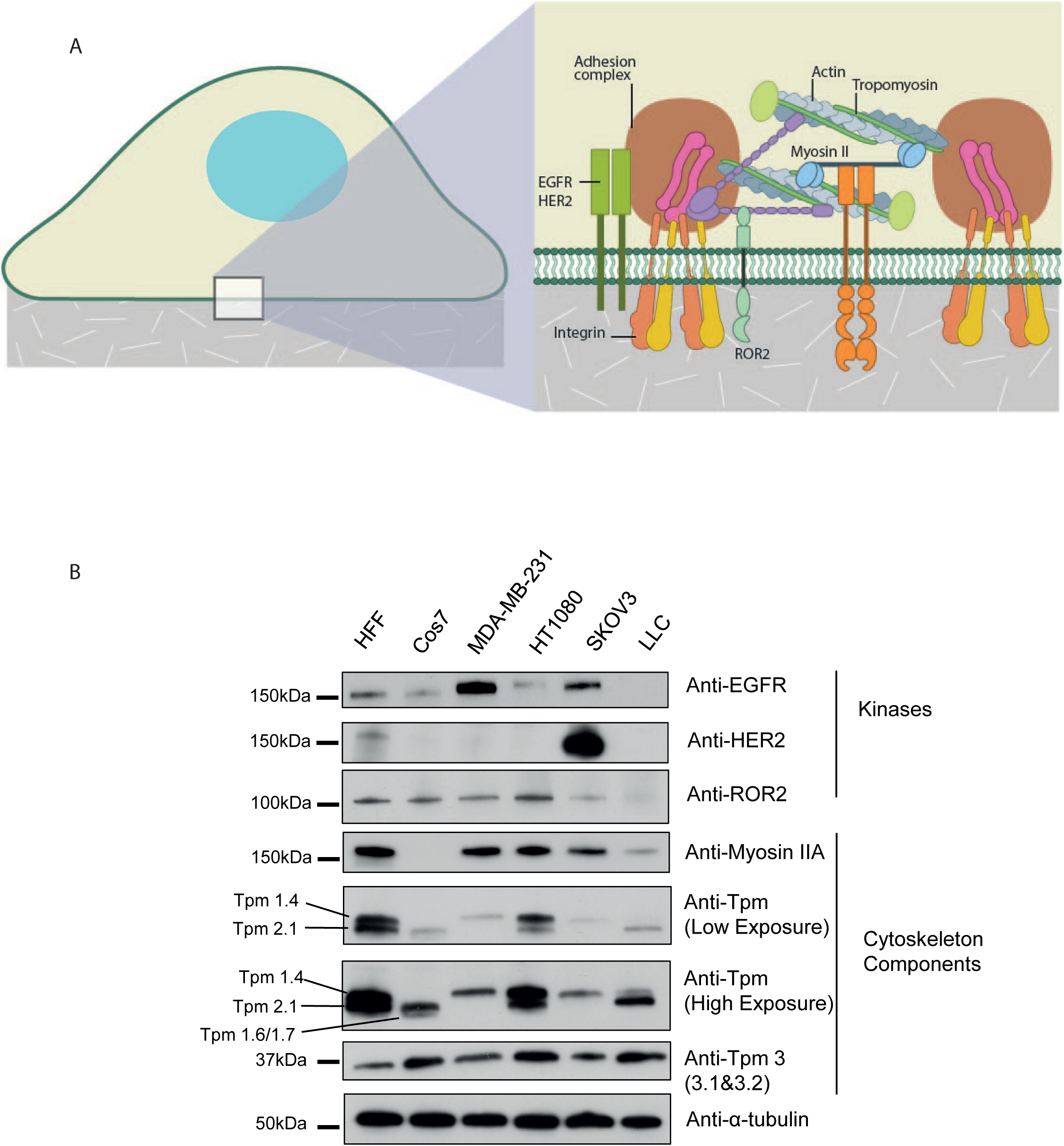
Transformed cells show altered expression of various mechanosensitive proteins. (A) A cartoon of the structure of a typical rigidity sensing contractile unit (CU) in HFF cells. (B) Western blots of the expression levels of different mechanosensitive components in HFF and different transformed cell lines.

### Cytoskeletal Proteins Restore Rigidity Sensing while Reciprocal Depletion Restores Cell Transformation

To address those questions, we first tested whether we could block transformed growth by simply restoring the missing cytoskeletal CU components for rigidity sensing. We selected Cos7 cells, which lacked myosin IIA, as the first candidate. After expression of EGFP-myosin IIA, Cos7 cells produced higher forces on both rigid and soft pillars (Figure 4 A). Also, the density of CUs increased dramatically in the presence of myosin IIA (Figure 4 B). We next examined whether restoration of CUs during initial spreading affected cell morphology and cell fate decision at later stages in the same environment. After 6 hours of spreading, Cos7 cells normally formed small focal adhesions of similar size on both soft (0.54±0.05 μm^2^) and rigid (0.58±0.08 μm^2^) flat PDMS surfaces (Figure 4 D and F). In contrast, Cos7-IIA (myosin IIA expressed Cos7 cells) generated larger focal adhesions (1.03±0.1 μm^2^) on rigid and smaller adhesions (0.65±0.03 μm^2^) on soft surfaces as commonly observed for normal fibroblast cell lines (Figure 4 C and F). In the colony growth assay on agar, Cos7-IIA cells did not survive after 7 days in culture whereas control Cos7 cells proliferated and formed colonies (Figure 4 G and H).

**Figure 4.**
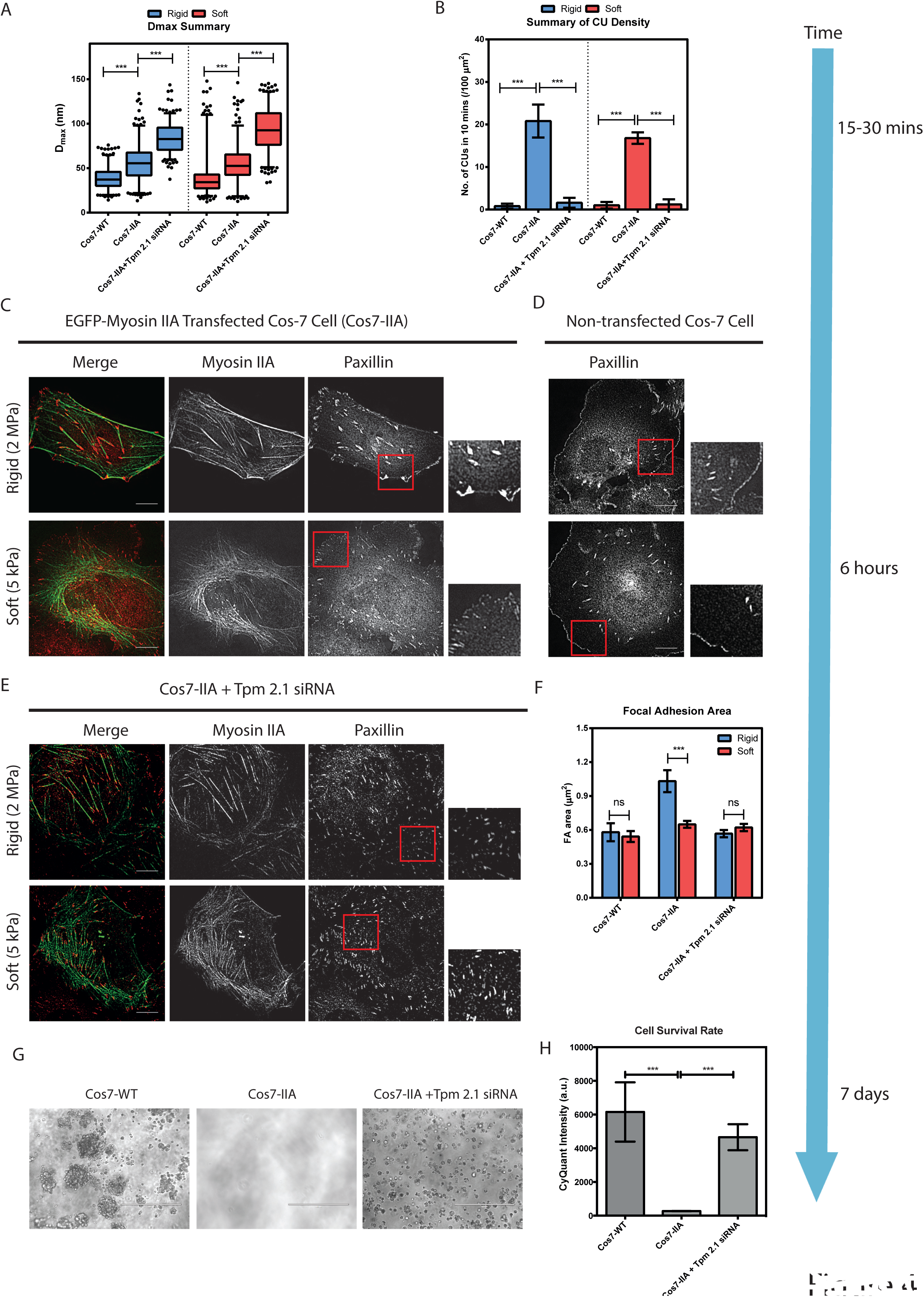
Re-expression of myosin IIA enables rigidity sensing and blocks transformed growth in Cos7 cells; Depleting Tpm 2.1 in Cos7-IIA cells restores transformed growth. (A) Box-and-whisker plots of the pillars’ maximum displacement values (D_max_) by Cos7-WT, Cos7-IIA and Tpm-siRNA-transfected Cos7-IIA cells on rigid (blue) and soft (red) pillars. Silencing Tpm 2.1 in Cos7-IIA cells increased the average force level on both types of pillars. (B) Bar graphs of average CU density per 10 minutes in Cos7-WT, Cos7-IIA and Tpm 2.1–siRNA-transfected Cos7-IIA cells on rigid (blue) and soft (red) pillars. (C, D and E) Paxillin images of Cos7 cells (C), EGFP-myosin IIA transfected Cos7 cells (Cos7-IIA)(D) and Tpm 2.1 depleted Cos7-IIA cells (E) fixed at 6 hours following seeding on rigid (2 MPa) or soft (5 kPa) fibronectin-coated PDMS surfaces. (Scale bar is 10 μm) (F) Mean single focal adhesion (FA) area of Cos7, Cos7-IIA and Tpm 2.1 silenced Cos7-IIA cells on rigid (blue) and soft (red) PDMS surface. (G and H) Soft agar assay indicating the growth of Cos7 and Tpm 2.1-knockdown Cos-7 IIA cells but not Cos7-IIA cells after 7-days culture. (Scale bar is 400 μm) (Error bars are SEMs. >500 pillars from >5 cells were analyzed in each condition; *** stands for p<0.001; ** stands for p<0.01; * stands for p<0.05)

To test if restoration of rigidity sensing activity in Cos7 cells was caused by myosin IIA and not by increasing the total amount of myosin in the cells, we next transfected myosin IIB in Cos7 cells. The Cos7-myosin IIB cells did not generate any CUs on pillar surfaces during initial spreading and they formed focal adhesions of similar size on both rigid (0.31±0.05 μm^2^) and soft (0.35±0.03 μm^2^) fibronectin-coated PDMS 6 hours after plating (Supplementary Figure 3). Thus, the re-expression of myosin IIA but not myosin IIB in Cos7 cells restored rigidity-sensing activity and blocked transformed growth.

We next asked whether reciprocal depletion of another CU component could reverse transformed growth of Cos7-IIA cells. Previously, depletion of Tpm 2.1 caused transformed growth in MCF-10A cells (Wolfenson et al., 2016). Upon siRNA depletion of endogenous Tpm 2.1 (Supplementary Figure 4 A), CU formation in Cos7-IIA cells was significantly suppressed (1.6 CUs /100 μm^2^ on rigid and 1.2 CUs/100 μm^2^ on soft pillars) (Figure 4 B). Despite the lack of CUs, Tpm 2.1-depleted Cos7-IIA cells produced larger displacements on both soft and rigid pillars, indicating that myosin-IIA was still active (Figure 4 A). Moreover, consistent with previous studies of MEF cells (Wolfenson et al., 2016), depletion of Tpm 2.1 in Cos7-IIA cells caused a significant decrease of FA size on glass surfaces (Figure 4 E and F). Further, Tpm 2.1-depleted Cos7-IIA cells survived on 2.3 kPa PAA gels without activating Caspase-3 cleavage after 3-days in culture (Supplementary figure 4 B and C). In the soft agar assay, visible colonies of Tpm 2.1-depleted Cos7-IIA cells formed (Figure 4 G and H). Thus, depletion of Tpm2.1 in Cos7-IIA cells blocked CU formation and restored transformed growth.

To investigate whether this connection between CU formation and transformed growth was cell line dependent, we examined the MDA-MB-231 cell line, which is a metastatic human breast cancer cell line that formed aggressive tumors in nude mice and was depleted of Tpm 2.1, (Freedman and Shin, 1974). Consistent with previous studies (Wolfenson et al., 2016), restoration of Tpm 2.1 in MDA-MB-231 cells decreased the pillar displacements (Figure 5 A) and restored CUs on both soft and rigid matrices (21.6 CUs/ 100 μm^2^ on rigid pillars and 14.8 CUs / 100 μm^2^ on soft pillars) (Figure 5 B). The other cell line that was missing Tpm 2.1 was SKOV3 and restoration of normal levels of Tpm 2.1 in those cells restored CUs as well (data not shown). The Tpm 2.1 transfected MDA-MB-231 cells (231-Tpm) distinguished between soft and rigid PDMS by generating larger focal adhesions and spreading to larger areas on rigid surfaces (Figure 5 C and F). Further, control MDA-MB-231 but not 231-Tpm cells formed colonies in soft agar culture after 7 days (Figure 5 G and H). In the reciprocal manner, depletion of endogenous myosin-IIA in 231-Tpm cells (Supplementary Figure 5 A) inhibited CU formation on both rigid and soft pillars by 5-fold (Figure 5 B) and decreased overall contractility (Figure 5 A). In addition, the myosin-IIA-depleted 231-Tpm cells showed a disruption of stress fibers and also a significant reduction of focal adhesion size on glass (Figure 5 E and F). Further, myosin-IIA-depleted 231-Tpm cells grew as well as wild type 231 cells on soft surfaces and the level of cleaved-Caspase-3 on soft surfaces also decreased 2-fold (Supplementary Figure 5 B and C). Additionally, after myosin-IIA silencing, 231-Tpm cells formed colonies in soft agar similar to MDA-MB-231 cells after 7 days (Figure 5 G and H). Thus, in the restored cell lines, depletion of other cytoskeletal components blocked CU formation and induced transformed cell growth. This indicated that rigidity sensing CU formation blocked anchorage-independent growth and loss of rigidity sensing enabled transformed growth in different cell backgrounds.

**Figure 5.**
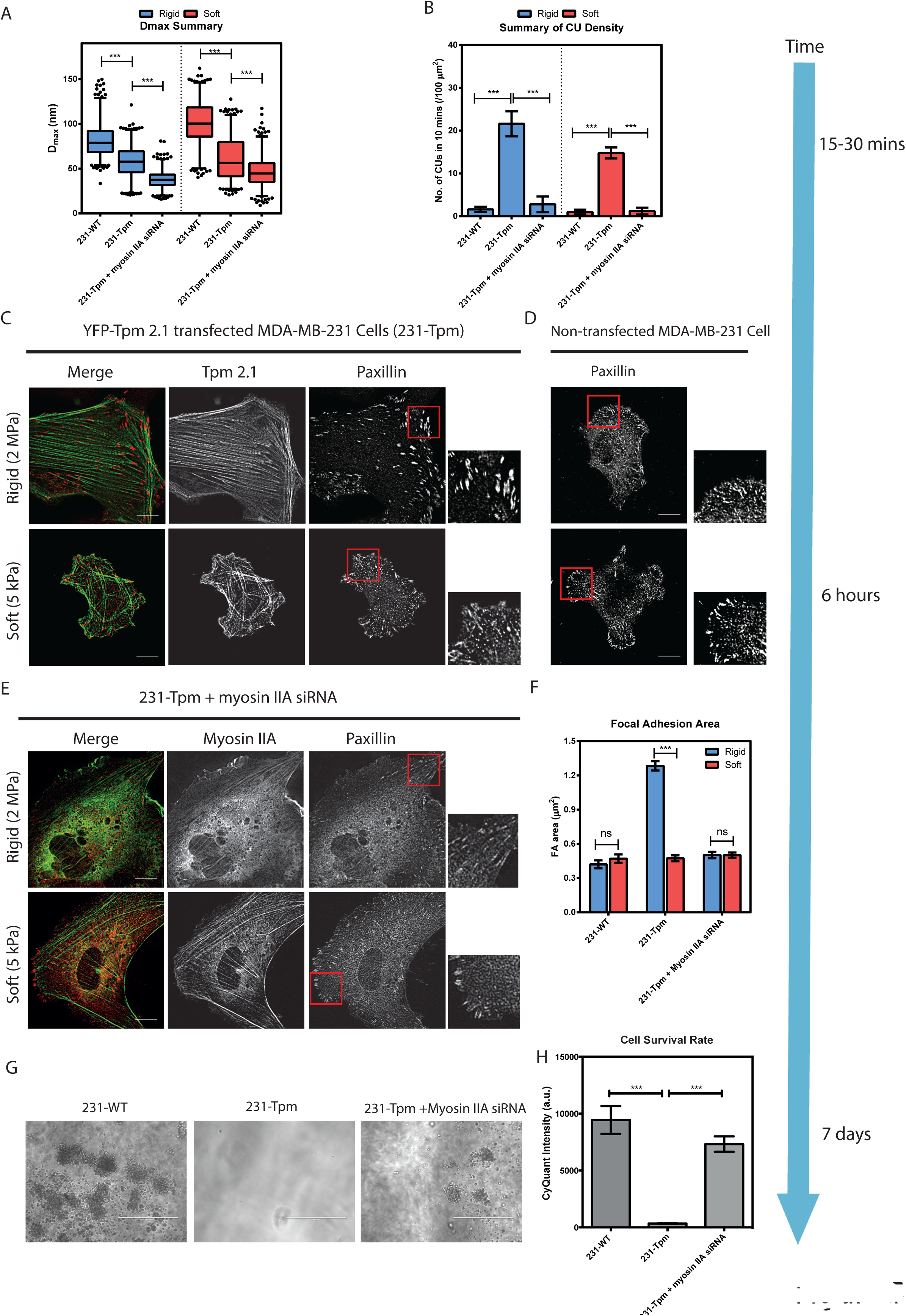
Tpm 2.1 expression in MDA-MB-231 cells enables rigidity sensing and blocks transformed growth; Depleting myosin IIA in 231-Tpm cells restores transformed growth. (A) Box-and-whisker plots of the pillars’ maximum displacement values (D_max_) by 231-WT, 231-Tpm and myosinIIA-siRNA-transfected 231-Tpm cells on rigid (blue) and soft (red) pillars. Silencing myosin IIA in 231-Tpm cells decreased the average force level on both types of pillars. (B) Bar graphs of average CU density per 10 minutes in different cells. (C, D and E) Paxillin images of MDA-MB-231 cells (C), YFP-Tpm 2.1 transfected MBA-MB-231 cells (231-Tpm)(D) and Myosin IIA depleted 231-Tpm cells (E) fixed at 6 hours following seeding on rigid (2 MPa) or soft (5 kPa) fibronectin-coated PDMS surface. (Scale bar is 10 μm) (F) Mean single focal adhesion (FA) area of 231, 231-Tpm and Myosin IIA silenced 231-Tpm cells on rigid (blue) and soft (red) PDMS surface. (G and H) Soft agar assay showing growth of MDA-MB-231 and Myosin-IIA-silenced 231-Tpm cells but not 231-Tpm cells after 7-days culture. (Scale bar is 400 μm) (Error bars are SEMs. >500 pillars from >5 cells were analyzed in each condition; *** stands for p<0.001; ** stands for p<0.01; * stands for p<0.05)

### A low molecular weight tropomyosin, Tpm 3, suppressed CU formation

Careful examination of the western blot screening data (Figure 3 B) raised a question about why HT1080 cells failed to generate CUs for rigidity sensing despite expressing the needed CU components. Notably, these cells also had high levels of another tropomyosin isoform, Tpm 3 (including Tpm 3.1 and Tpm 3.2), which was highly expressed in many cancers (Stehn et al., 2013). To determine if Tpm 3 suppressed CU formation, we silenced the endogenous Tpm 3 in HT1080 cells by siRNA (Figure 6 A). As shown in Figure 6 B, after Tpm 3 knockdown, the pillar displacements were significantly lower than those of the control group on both rigid and soft pillars.

**Figure 6.**
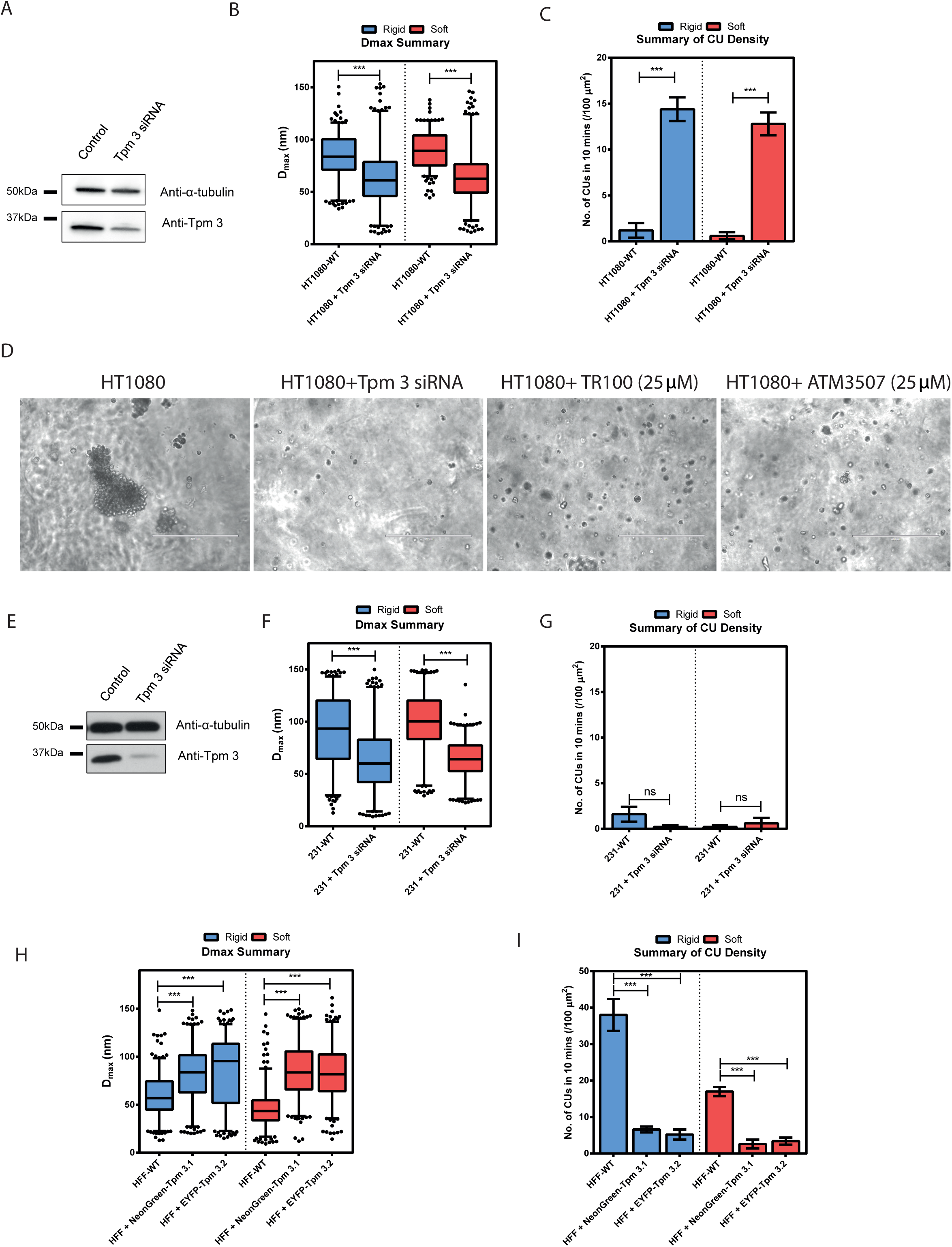
Tpm 3 (3.1 and 3.2) levels affect cell rigidity sensing. (A)Western blot showing Tpm 3 (Tpm 3.1 and 3.2) levels in HT1080 cells treated with scramble or anti-Tpm 3 siRNA. (B) Box-and-whisker plots of the pillars’ maximum displacement values (D_max_) control or Tpm 3 depleted HT1080 cells on rigid (blue) and soft (red) pillars. (C) Average CU density per 10 mins for control and Tpm 3 depleted HT1080 cells on two pillar substrates of different rigidities. (D) Soft agar assay showing growth of control HT1080 cells but not of Tpm 3 depleted or cells treated with Tpm 3 inhibitors (TR100 and ATM 3507). (Scale bar is 400 μm) (E) Western blot showing Tpm 3 levels in MDA-MB-231 cells treated with scrambled or anti-Tpm 3 siRNA. (F) Box-and-whisker plots of the pillars’ maximum displacement values (D_max_) control or Tpm 3 depleted MDA-MB-231 cells on rigid (blue) and soft (red) pillars. (G) Average CU density per 10 mins for control and Tpm 3 silenced MDA-MB-231 cells on two pillar substrates of different rigidities. (H) Box-and-whisker plots of the pillars’ maximum displacement values (Dmax) generated by control HFF cells, NeonGreen-Tpm 3.1 overexpressed or EYFP-Tpm 3.2 overexpressed HFF cells on rigid (blue) and soft (red) pillars. (I) Average CU density per 10 min of control, NeonGreen-Tpm 3.1 or EYFP-Tpm 3.2 transfected HFF cells on two pillar substrates of different rigidities. (Error bars are SEMs. >500 pillars from >5 cells were analyzed in each condition; *** stands for p<0.001; ** stands for p<0.01; * stands for p<0.05)

In addition, the Tpm 3-depleted HT1080 cells generated 14.4 CUs/100 μm^2^ on rigid pillars and 10.8 CUs/100 μm^2^ on soft pillars, approximately 5-fold higher than control. Moreover, the depletion of Tpm 3 or inhibition of Tpm 3 assembly on actin filaments by TR100 or ATM 3507 (Stehn et al., 2013) decreased the size and number of colonies that HT1080 cells formed in soft agar after 7-days in culture (Figure 6 D). Thus, these results supported the idea that the high level of Tpm 3 protein in HT1080 cells suppressed CU formation and stimulated transformed growth.

To check if Tpm 3 depletion would activate CU formation even in the absence of Tpm 2.1, we knocked down Tpm 3 by siRNA in MDA-MB-231 cells which lacked endogenous Tpm 2.1 (Figure 6 E). Although Tpm 3 depletion caused a decrease of force level in MDA-MB-231s (Figure 6 F), it failed to increase CU density on both rigid and soft surfaces (Figure 6 G). This indicated that Tpm 2.1 expression was necessary for CU formation and cell rigidity sensing.

Decreased expression of high molecular weight tropomyosins (including Tpm 2.1) and increased expression of low molecular weight tropomyosins (including Tpm 3.1 and Tpm 3.2) were reported in cells transformed by various oncogenes, carcinogens and viruses (Helfman et al., 2008). The fact that silencing Tpm 3 in Tpm 2.1-expressing transformed cells increased the number of CUs during early spreading suggested that there was a competition between Tpm 2.1 and Tpm 3 (Tpm 3.1 and Tpm 3.2) during CU formation. To test this hypothesis, we asked whether overexpression of Tpm 3 (Tpm 3.1 or Tpm 3.2) in a normal fibroblast cell line would suppress cell rigidity sensing. To that end, HFF cells expressing high levels of NeonGreen-Tpm 3.1 or EYFP-Tpm 3.2 were analyzed on pillar surfaces. Forces produced by these cells were significantly higher than wild-type HFF cells on both rigid and soft surfaces (Figure 6 H). Conversely, the CU density was decreased (Figure 6 I). This indicated that high expression levels of Tpm 3.1 or Tpm 3.2 inhibited CU formation, potentially through competing with Tpm 2.1.

**Figure 7.**
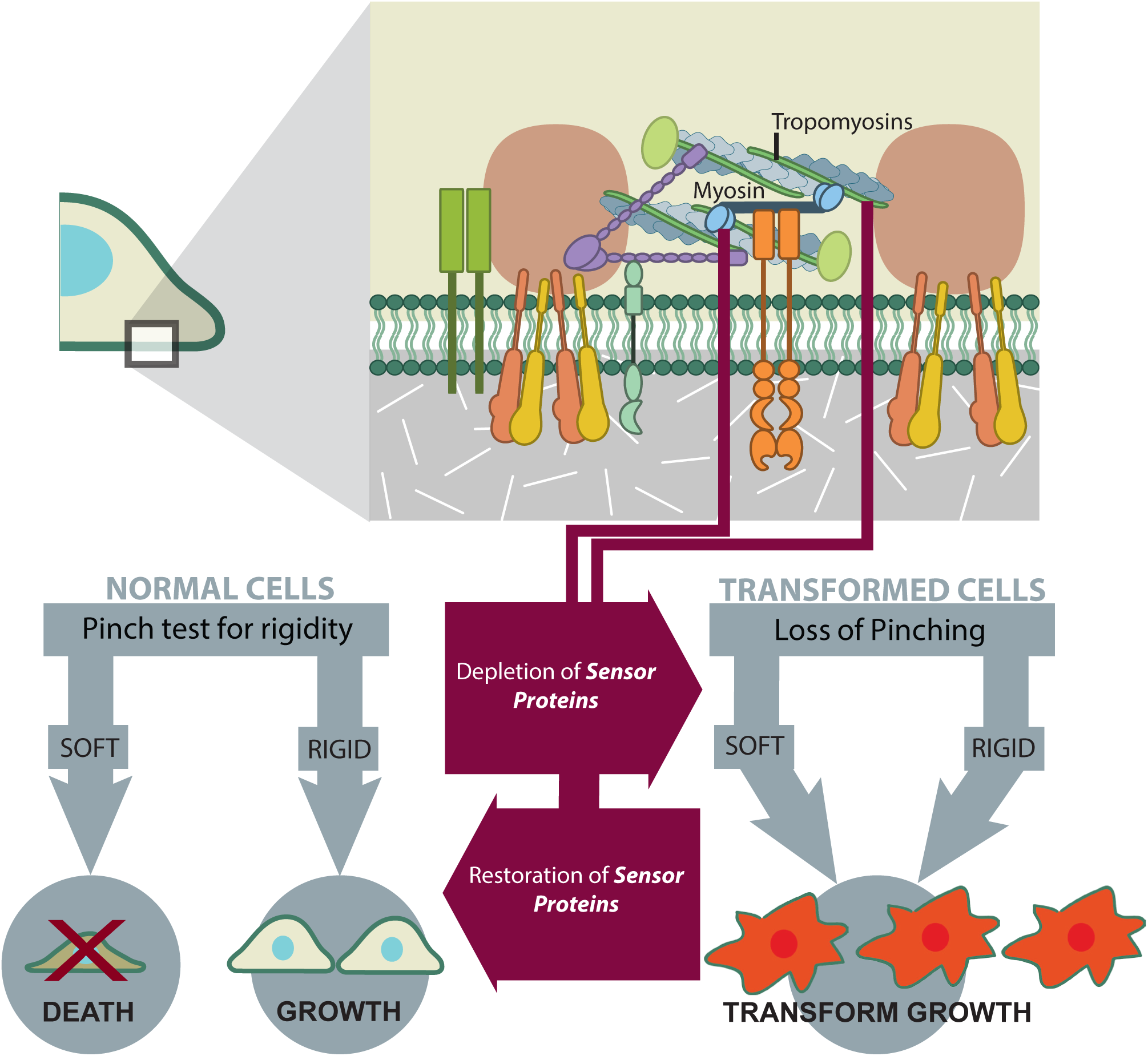
Proposed Model of the Role that the loss of Rigidity Sensing Contractile Units Plays in Transformed Growth.

To further examine the behavior of Tpm 2.1 in the Tpm 3.1-overexpressing cells, we fixed and stained the endogenous Tpm 2.1 in HFFs with or without NeonGreen-Tpm 3.1 overexpression after 15 mins spreading on pillars. Consistent with previous observations (Wolfenson et al., 2016), Tpm 2.1 concentrated at cell edges in control HFFs. However, this peripheral localization of Tpm 2.1 disappeared in Tpm 3.1 overexpressing cells (Supplementary Figure 6). Thus, we suggest that high levels of Tpm 3 inhibited cell rigidity sensing CU formation by competing with endogenous Tpm 2.1 in normal fibroblasts.

## Discussion

Based upon the current and related findings, transformed cell growth in cancer cells correlates with the inability of cells to sense substrate rigidity, regardless of whether rigidity sensing is inhibited by depletion of myosin IIA, Tpm 2.1 or α-actinin 4, as well as overexpression of Tpm 3. In widely different cell backgrounds of cancer cells from four different tissues and African green monkey kidney cells transformed by SV-40, the unbalanced expression of a distinct set of cytoskeletal proteins can decrease rigidity sensing contractile unit (CU) formation, which results in transformed growth. When rigidity sensing CUs were restored in these highly different transformed cells, there was rigidity-dependence of growth, i.e., growth on rigid surfaces and activation of Caspase-3-dependent *anoikis* on soft matrices. In the initial characterization of CUs, the major cytoskeletal components, Tpm 2.1, myosin-IIA, and α-actinin 4, were found to be necessary components for CU formation (Meacci et al., 2016; Wolfenson et al., 2016). In these studies, overexpression of another cytoskeletal protein, Tpm 3 (including Tpm 3.1 and Tpm 3.2), causes CU disruption. These findings strongly support the hypothesis that transformed growth in cancer cells requires the depletion of rigidity sensing.

In the general context of cancer, there is a strong link between the cytoskeletal proteins involved in rigidity sensing and the disease. Tpm 2.1, Myosin IIA and α-actinin 4 are reported to be tumor suppressors and Tpm 3 is a tumor promoter. The strongest case is for Tpm 2.1 that is down-regulated in a wide variety of transformed cell lines (Bhattacharya et al., 1990; Hendricks and Weintraub, 1981; Masuda et al., 1996; Shah et al., 1998). Overexpression of miR-21, a microRNA that targets the Tpm 2.1 RNA (Zhu et al., 2007), is observed in breast tumors and correlates with the severity of the disease (Si et al., 2006). Further, reduction of miR-21 induces glioma cell apoptosis through Caspase pathways (Zhou et al., 2010). In an *in vivo* RNAi screen, myosin-IIA has been identified as a tumor suppressor of squamous cell carcinomas (Schramek et al., 2014). α-actinin 4, as well, has been recognized as a tumor suppressor in cases of neuroblastoma and lung cancers (Menez et al., 2004; Nikolopoulos et al., 2000). In contrast, Tpm 3 is responsible for metastatic melanoma motility regulation (Gunning et al., 2008) and expression levels are elevated in many different cancer lines (Stehn et al., 2013). Thus, cancer cells that are characterized by transformed growth on soft surfaces, have altered levels of various cytoskeletal proteins that correlate with the loss of local CUs involved in rigidity sensing but do not correlate with changes in tyrosine kinase levels.

One of the major pathways to induce transformation is to express mutant Ras, which is normally activated by a variety of RTKs that participate in cell proliferation, transformation, and regulation of differentiation (Yamamoto et al., 1999). For example, MDA-MB-231 cells carry a K-Ras mutation (Scholl et al., 2009). In previous studies, mutant Ras isoforms caused a depletion of tropomyosin 2.1 possibly through the up-regulation of miR-21 (Frezzetti et al., 2011). In terms of other ways of transforming cells, Cos7 is an SV40 transformed cell line. SV40 transformation in MEF cells increases Tpm 3 but not Tpm 2.1 expression levels (Coombes et al., 2015). As we show here, overexpression of Tpm 3 decreases the number of CUs by competing with Tpm 2.1. Thus, classical ways of transformation normally alter different cytoskeletal protein expression levels and therefore cause depletion of CUs. This further supports the relationship between rigidity sensing CU formation and cell transformation.

An additional connection between cancer and the local CUs comes from the role that tyrosine kinases play in rigidity sensing. Previous studies show that rigidity sensing requires the action of the Src family kinases (SFKs). Knocking out the upstream activator, RPTPα, or the three Src kinases, Src, Yes, and Fyn (SYF cells) blocks the ability of those cells to sense substrate rigidity and also enables growth on soft surfaces (Jiang et al., 2006; Sawada et al., 2006). This is consistent with our recent study that showed that ErbB family members (EGFR and HER2) are recruited to adhesion sites by SFKs on rigid surfaces and catalyze the formation of CUs in early cell spreading on rigid surfaces (Saxena et al., 2017). These findings indicate that RTKs involved in cancer and in epithelial-to-mesenchymal transition also play important roles in rigidity sensing regulation.

There were dramatic differences in the level of expression of the various tyrosine kinases between the different transformed cell lines. This does not mean that RTKs do not contribute to the potential of cancer cells to grow. Rather, it seems that cytoskeleton changes altered the mechanical signals that cells collected from their microenvironment and RTKs expanded those signals for further cell fate decision making. Thus, the role of RTKs in transitions between different cell states is more complicated.

Transformation involves a major change in cell behavior that can be described as a change in cell state. It is therefore surprising that low levels of Tpm 2.1, myosin IIA or α-actinin 4 can cause transformation in a variety of cell backgrounds. When myosin IIA is present, the transformed cells develop very high forces on the 500 nm pillars but in the absence of myosin IIA there is a much lower level of force. MDA-MB-231 cells show rapid transformed growth without myosin IIA, indicating that the loss of rigidity sensing not the level of force at these initial stages activates transformed growth. Myosin IIA is normally present in most cancer cells and they then develop very high forces on substrates (Mierke et al., 2011), which may aid growth in an *in vivo* context.

Severe cancers have many features that are important for tumor growth; however, the depletion of rigidity sensing modules that causes transformation enables them to escape from the *anoikis* pathways and to adopt the transformed phenotype. It is remarkable that re-introduction of the missing CU components in transformed cancer cell lines successfully rebuilds the rigidity sensing process irrespective of the tissue of origin. Restoring the ability to correctly sense rigidity causes many transformed cancer cells to die on soft surfaces without further manipulations. This provides a different view of blocking transformed growth. Namely, a functional rigidity sensor activates apoptosis pathways on soft surfaces and the depletion of those sensors is often sufficient for cell growth. Although there may be other mechanisms for causing growth on compliant matrices, the loss of the rigidity-sensing modules is a robust mechanism. As modular sensory units in self-driving cars are needed to stop them when an object is in their path, the modular rigidity-sensing units are needed for cells to know when the matrix is soft and growth should stop. The observation that depleting various components of rigidity-sensing modules depletes rigidity-sensing indicates that like many complex sensory modules, the rigidity-sensing contractile units require many different proteins. It is easy, therefore, to understand how many different mutations or alterations of cells could result in transformation by the loss of a single sensory machine and why cancer is such a difficult disease to treat.

## Material and Methods

### Cell culture and transfection

HFF cells (ATCC), Cos7 cells (ATCC), MDA-MB-231 cells (gift from Dr. Jay Groves, MBI, NUS), HT1080 (ATCC), SKOV3 cells (gift from Dr. Ruby Huang, CSI, NUS) and LLC cells (ATCC) were cultured in DMEM with high glucose supplemented with 10% FBS and 1mM sodium pyruvate. Cells were transfected with DNA plasmids using lipofectamine 2000 (Invitrogen) or by Neon electroporator system (Life Technologies) according to the manufacturer’s instructions. Expression vectors encoding the following fluorescent fusion proteins were used: Tpm 2.1-YFP, NeonGreen-Tpm 3.1 (also named as Tm 5NM1), EYFP-Tpm 3.2 (also named as Tm 5NM2) (gifts from Dr. Peter Gunnning), MyosinIIA-EGFP and Emerald-myosin-IIB (gift from Dr. Michael W. Davidson group).

### Transfection of siRNA and immunoblotting

Cells were seeded into a 6-well dish on day 0 and transfected with 25 μM myosin-IIA siRNA (Dharmacon), Tpm 2.1 siRNA (Qiagen) or Tpm 3 siRNA (Dharmacon) using lipofectamine RNAiMAX (Invitrogen) on day 1. Control cells were transfected with scrambled control siRNA (Dharmacon). Transfected cells were lysed in RIPA buffer (Sigma) and proteins extracted were separated by 4-20% SDS-polyacrylamide gel (Bio-rad) and transferred to PVDF membranes (Bio-rad) at 75V for 2 hours. Membranes were incubated with appropriate primary antibodies at 4°C overnight: anti-myosin-IIA (Sigma, dilution 1:1000), anti-Tpm 2.1 (Abcam, dilution 1:1000), anti-TM311 (Sigma, dilution 1:1000), anti-TM γ9d (gift from Dr. Peter Gunning, dilution 1:1000), anti-EGFR (CST, dilution 1:1000), anti-HER2 (CST, dilution 1:1000), anti-ROR2 (CST, dilution 1:1000) and anti-α-tubulin (Sigma, dilution 1:3000). The primary antibody binding was processed for ECL detection (Thermo Fisher Scientific) with appropriate HRP-conjugated secondary antibodies (Bio-rad).

### Pillar fabrication, video microscopy and force traction measurements

Molds for making PDMS pillars were fabricated as described before (Saxena et al., 2017). 0.1g of PDMS (mixed at 10:1, Sylgard 184; Dow Corning) was poured onto the silicon mold and then flipped onto a plasma-cleaned glass bottom dish (ibdi). The sample was pressed by an 8g weight, cured at 80°C for 3 hours to reach a Young’s modulus of 2 MPa and was de-molded while immersed in 99.5% isopropanol. Pillars were washed with PBS for 5 times before coating with 10 μg/ml fibronectin (Roche) for cell seeding. Time-lapse imaging and traction force measurements were performed as explained before (Saxena et al., 2017; Yang et al., 2016).

### PAA gel preparation

Glass bottom dishes (Iwaki) were silanized using 1.2% 3-methacryloxypropyltrimethoxysilane (Shin-Etsu Chemical, Tokyo, Japan) in 100% Methanol for 1 hour at room temperature. 2.3 kPa Acrylamide gel was prepared as previously described (Nakazawa et al., 2016). Gel surfaces were treated with sulfo-SANPAN (Thermo Fisher Scientific) and exposed under UV for 5 mins before coating with 10 μg/ml fibronectin for cell culture.

### Fluorescence microscopy

Cells were fixed with 4% paraformaldehyde in PBS at 37°C for 15 mins and permeabilized with 0.2% TX-100 for 10 mins at room temperature. Samples were blocked with 1% bovine serum albumin (BSA) in PBS for 1h at room temperature, incubated with primary antibodies for paxillin (BD, 1:200) or Cleaved-Caspase-3 (CST, 1:200) at 4 °C overnight and then incubated with secondary antibodies (Molecular Probes) for 1h at room temperature. Fluorescence images were acquired using a spinning-disc confocal microscope (PerkinElmer Ultraview VoX) attached to an Olympus IX81 inverted microscope body.

### Soft agar assay

The soft agar assay was performed using the Cell Transformation Assays, Standard Soft Agar Kits from Cell Biolabs according to manufacturer’s instructions.

### Statistical analysis

Prism (GraphPad Software) and Matlab (Math Works) were used for data analysis and graph plotting. Analyses of significant difference levels were carried out using ANOVA test (for more than 2 experimental groups) or Student’s t-test.

## Acknowledgements

The authors thank JP Thiery, P Gunning, NC Gauthier, RYJ Huang and J Kadrmas for the critical reading of the manuscript. We thank all the members of Sheetz lab and Bershadsky lab for their kind help. This research was supported by funding to the MBI, National University of Singapore. B.Y was supported by NUS grant “Activation of Apotosis by Soft Surfaces” (R-714-000-112-133). H.W. is a David and Inez Myers Career Advancement Chair in Life Sciences fellow. M. P. S is supported by NIH grants, NUS grants and Mechanobiology Institute, National University of Singapore.

## Author Contributions

B.Y. and M.P.S. conceived the study and designed the experiments; B.Y. and N.N. performed the experiments; S.L. wrote Matlab codes for data analysis; B.Y. analyzed the data; J.H. provided fabrication molds; B.Y. H.W. and M.P.S wrote and prepared the manuscript.

---

**Supplementary Figure 1** (A) Displacement vs. time plot of one pillar that was not touched by the cell (reference pillar) during imaging. (B) Box and whiskers plots of pillar maximum displacement summary from 200 reference pillars on rigid (blue) and soft (red) pillars. There is no significant difference between the system noise levels on rigid or soft pillar substrates.

**Supplementary Figure 2** (A) Protein expression heat map summary of various mechanosensitive proteins in various transformed cell lines.

**Supplementary Figure 3** (A) Myosin-IIB and paxillin images of Cos7 cells transfected with Emerald-myosin-IIB (Cos7-IIB) on soft and rigid PDMS surfaces. (Scale bar is 10 μm) (B) Bar graphs of the mean values of single focal adhesion areas of Cos7-IIB cells on rigid and soft surfaces (Error bars are SEMs. >200 focal adhesions from >5 cells were analyzed in each condition).

**Supplementary Figure 4** (A) Western Blot showing Tpm 2.1 levels in Cos7 cells treated with scramble or anti-Tpm 2.1 siRNA. (B) Caspase-3 and DAPI staining images of Cos7-WT, Cos7-IIA and Tpm 2.1 depleted Cos7-IIA cells after 3 days culturing on 2.3 kPa fibronectin coated PAA gel surfaces. (Scale bar is 50 μm) (C) Average Caspase-3 intensity level measurements of cells in different conditions (Error bars are SEMs; >30 cells were analyzed in each case; *** stands for p<0.001; ** stands for p<0.01; * stands for p<0.05).

**Supplementary Figure 5** (A) Western blot showing Tpm 3 levels in HT1080 cells treated with scramble or anti-Tpm 3 siRNA. (B) Caspase-3 and DAPI staining images of 231-WT, 231-Tpm and myosin IIA depleted 231-Tpm cells after 3 days culturing on 2.3 kPa fibronectin coated PAA gel surfaces. (Scale bar is 50 μm) (C) Average Caspase-3 intensity level measurements of cells in different conditions (Error bars are SEMs; >30 cells were analyzed in each case; *** stands for p<0.001; ** stands for p<0.01; * stands for p<0.05).

**Supplementary Figure 6** (A) Fluorescence images of NeonGreen-Tpm 3.1 and endogenous Tpm 2.1 in HFF cells spreading on rigid pillar surfaces. (B) Average total endogenous Tpm 2.1 intensity in control and Tpm 3.1 overexpressed HFF cells. (C) Average endogenous Tpm 2.1 intensity near cell periphery in control and Tpm 3.1 overexpressed HFF cells. (D) Cell aspect ratio of control and Tpm 3.1 overexpressed HFF cells after 15 mins spreading on rigid pillar surfaces. (*** stands for p<0.001; ** stands for p<0.01; * stands for p<0.05)

